# Revealing multiscale architecture of the intact human spine using hierarchical phase-contrast tomography

**DOI:** 10.64898/2026.10.23.760441

**Authors:** Aikta Sharma, Isla C. Henderson, Alissa L. Parmenter, Joseph Brunet, Christian Liebsch, Ann-Kathrin Greiner-Perth, David Stansby, Jishizhan Chen, Yang Zhou, Jingwen Liu, Joanna Purzycka, Theresa Urban, Hector Dejea, Aurélien Delphin, Laurent Lamalle, Andrew A. Pitsillides, Timothy J. P. Bray, Hans-Joachim Wilke, Alexandre Bellier, Paul Tafforeau, Claire L. Walsh, Peter D. Lee

## Abstract

**Background:** CT and MRI provide detailed clinical assessment of the human spine; however, their millimeter-scale spatial resolution limits direct visualization of tissue microarchitecture and interfaces. Hierarchical phase-contrast tomography (HiP-CT) enables *ex vivo* non-destructive, micron-resolution imaging of intact human organs without exogenous contrast agents, extending anatomical assessment to spatial scales inaccessible to conventional clinical imaging.

**Purpose:** To evaluate the capability of HiP-CT to visualize macro- to microanatomy of an intact adult human spine *ex vivo*.

**Materials and Methods:** An intact spine (C1-L5) from a 62-year-old male donor without known skeletal disease was formalin-fixed and imaged by HiP-CT at the European Synchrotron. Whole-spine imaging was performed using 16.26 µm isotropic voxels, with selected regions of interest imaged using 2.2 µm voxels. *Ex vivo* MRI was performed for anatomical comparison. Three-dimensional segmentation and quantitative analyses included intervertebral disc (IVD) grading, annulus fibrosus fiber tracing, and vertebral trabecular micro-morphometry.

**Results:** HiP-CT enabled high spatial-resolution, multiscale imaging of the intact human spine, simultaneously visualizing vertebral bone and endplate, IVDs, spinal cord and meninges, nerve roots and ganglia, vasculature, ligaments, and musculature. Three-dimensional segmentation and virtual sectioning mapped neurovascular anatomy and vertebral vascular supply, without exogenous contrast enhancement, while preserving their relationships with surrounding tissues. Whole-spine images enabled quantitative analysis of annulus fibrosus fiber organization and vertebral trabecular microarchitecture, permitting the detection of focal posterolateral fiber disruption at the L4-L5 IVD and regional differences in trabecular architecture between cervical and lumbar vertebral bodies. Local tomography resolved annulus fibrosus lamellae, vertebral endplate Haversian systems, and facet joint hyaline cartilage at micron resolution.

**Conclusions:** HiP-CT enables non-destructive, multiscale three-dimensional imaging of the intact human spine while preserving tissue relationships across spatial scales and tissue boundaries. HiP-CT images could provide an *ex vivo* microanatomical reference for future correlation with clinical imaging investigations of spinal disease and in the discovery and validation of imaging biomarkers.

**Key results:**

- Synchrotron-based hierarchical phase-contrast tomography enables non-destructive, multiscale three-dimensional imaging of an intact adult human spine using 16.26 µm isotropic voxels, resolving both mineralized and non-mineralized tissue.
- Local tomography using 2.2 µm isotropic voxels reveals additional microstructural detail without exogenous contrast agents or sectioning.
- Whole-spine imaging enables quantitative characterization of intervertebral disc fiber architecture and vertebral trabecular morphometry, alongside visualization of neurovascular anatomy while preserving spatial relationships across the intact specimen.

**Summary statement:** Hierarchical phase-contrast tomography enables non-destructive multiscale imaging of the intact human spine, resolving microarchitecture across tissues and interfaces and providing a basis for bridging clinical imaging with underlying three-dimensional microanatomy.

## Introduction

The human spine is a complex, hierarchically organized organ in which vertebrae, intervertebral discs (IVDs), neural and vascular tissues are structurally and functionally integrated across spatial scales. Understanding their three-dimensional architecture and spatial relationships is fundamental to defining structure-function relationships that underpin spinal health and disease.

CT and MRI are commonly used modalities for *in vivo* assessment of spinal anatomy. CT is widely used for fracture, trauma, postoperative and degenerative assessment (1-4), whereas MRI offers superior soft tissue contrast for assessing IVDs, the spinal cord, nerve roots and paraspinal tissues, particularly in suspected malignancy, infection, or neurological compromise (5-7). However, spatial resolution is limited to the millimeter-scale (8, 9), and despite advances in photon-counting CT (10) and ultra-high field MRI (11), comprehensive three-dimensional visualization of spinal tissue microarchitectures remains challenging. Although quantitative MRI techniques, including relaxometry, susceptibility mapping and diffusion imaging, yield additional *in vivo* information on spinal tissue composition and organization (12-15), they offer only indirect measures of microstructure and remain subject to acquisition time, signal-to-noise, and field inhomogeneity constraints.

*Ex vivo* imaging provides an opportunity to directly interrogate tissue architecture at substantially finer spatial scales. Micro-CT achieves micron-scale resolution in mineralized tissues but typically requires exogenous contrast enhancement for soft tissue visualization and is limited by specimen size and field of view (FOV) (16-18). Histology, electron microscopy, and confocal microscopy, in contrast, achieve cellular-to-subcellular detail yet necessitate destructive sample preparation and thus do not preserve three-dimensional relationships across intact spinal structures (19-22). Synchrotron phase-contrast imaging can overcome these limitations by enhancing soft tissue visibility, as demonstrated in human spinal cord and column specimens (23). Hierarchical phase-contrast tomography (HiP-CT) extends this approach to rapid, non-destructive, multiscale imaging of intact human organs at approximately 20 µm isotropic voxel size for whole-organ imaging and 1-6 µm voxels for local tomography, offering new insights into physiology and pathophysiology across a range of organ systems (24-29). However, HiP-CT has not yet been applied to spinal imaging.

In this proof-of-concept study, we evaluated whether HiP-CT could be used to image a single intact adult human spine (C1-L5) across macro- to microscopic scales, using clinical MRI as a comparator. We assessed its ability to resolve neurovascular anatomy and characterize annulus fibrosus fiber organization alongside vertebral trabecular microarchitecture (Figure 1) within their native anatomical contexts, and whether HiP-CT images could constitute a three-dimensional microanatomical reference for relating clinical imaging findings to underlying tissue microarchitecture.

**Figure 1:**
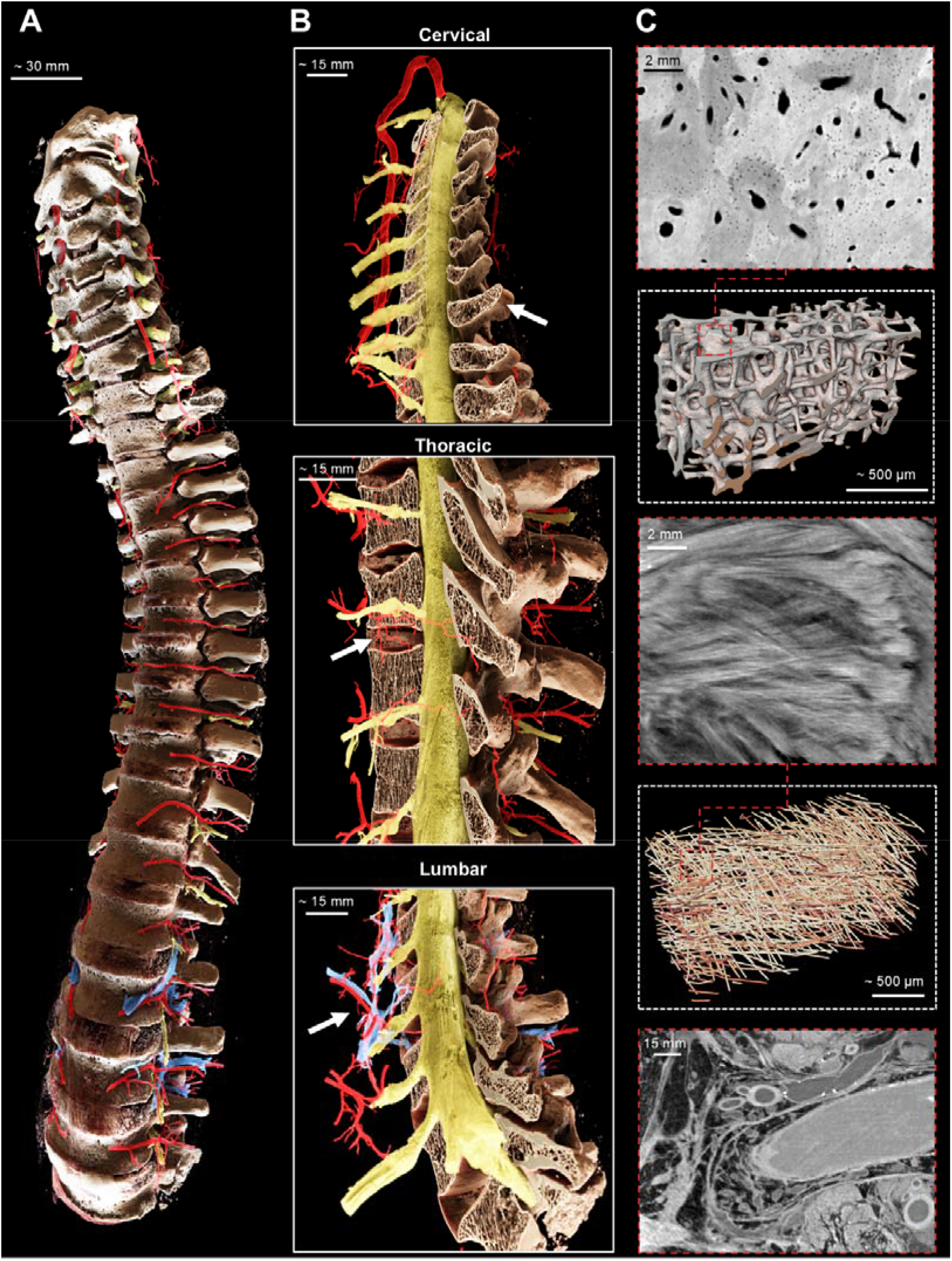
Multiscale HiP-CT visualization of the anatomy of an intact human spine. Three-dimensional anatomical rendering of the intact human spine showing the vertebral column (cream), spinal cord and nerve roots (yellow) and associated arterial (red) and venous vasculature (blue; A). Zoomed sagittal sectioned views of representative cervical (top), thoracic (middle), and lumbar (bottom) regions (B) show osseous, neural, and vascular anatomy across the length of the spine. Representative high-resolution HiP-CT images (red hatched boxes) and selected three-dimensional renderings of regions indicated by white arrows in B illustrate hierarchical tissue architectures resolved across spatial scales. These include cervical vertebral bone microarchitecture, with intracortical vascular canals visible within Haversian systems and trabecular organization (C, top); the lamellar architecture of the thoracic IVD annulus fibrosus and its three-dimensional fiber organization (C; middle); soft tissue and vascular structures adjacent to the spine (C, bottom).

## Materials and Methods

Additional methodological and technical details are provided in the Supplementary Material (Figure S1; Tables S1 and S2).

### Specimen description

A complete spine was obtained from a single body donated to the Laboratoire d’Anatomie des Alpes Françaises (LADAF) in November 2023, in accordance with French legislation governing body donation. The French Ministry of Health approved subsequent transport and imaging protocols. The donor was a 62-year-old White male with no known skeletal disorders (Table 1), and the sample and associated donor information were anonymized prior to HiP-CT imaging. Following dissection, the intact spine (C1-L5) including the major vessels, nerves, spinal cord, vertebrae, facet joints, and IVD measured 62 x 13 x 17 cm in the cranio-caudal, medio-lateral, and anterior-posterior dimensions, respectively, and was fixed by immersion in 4% (v/v) neutral buffered formalin prior to imaging.

**Table 1:**
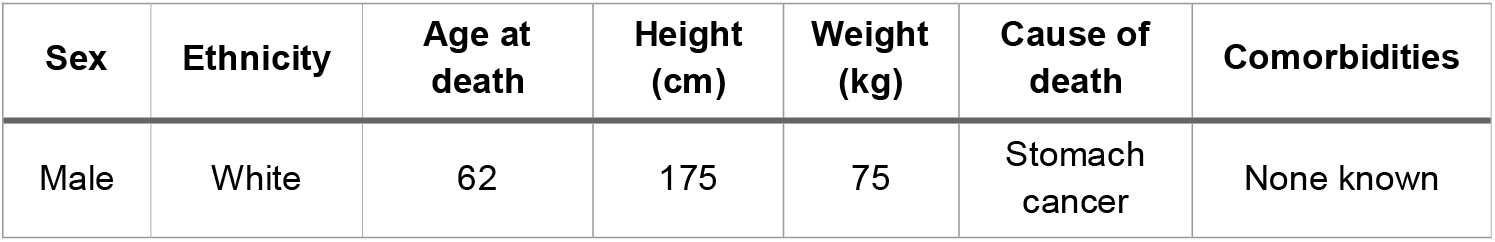
Medical information of the organ donor.

| Sex | Ethnicity | Age at death | Height (cm) | Weight (kg) | Cause of death | Comorbidities |
| --- | --- | --- | --- | --- | --- | --- |
| Male | White | 62 | 175 | 75 | Stomach cancer | None known |

### MR Image acquisition

*Ex vivo* MRI was performed at IRMaGe at Grenoble Central University Hospital, France using a 3 T Achieva dStream (Philips Medical Systems, Netherlands) approximately three weeks prior to HiP-CT. Sagittal two-dimensional T1-weighted (T1w), T2-weighted (T2w), T2*-weighted (T2*w) and T2-weighted short tau inversion recovery (T2-STIR) sequences were acquired across three contiguous anatomical stations to accommodate the length of the specimen and stitched to generate whole-spine images.

### HiP-CT Image acquisition and reconstruction

The intact spine was positioned in an anatomically representative orientation, stabilized in agar prepared in 4% neutral buffered formalin (v/v) and subjected to in-line degassing as previously described (24). HiP-CT imaging was performed at the BM18 beamline (European Synchrotron, Grenoble, France) using a filtered, polychromatic X-ray beam with average energies ranging from 87-160 keV (Figure S1). The whole-organ overview scan was acquired using continuous helical tomography with an isotropic voxel size of 16.26 µm. Local tomographic scans of 8 volumes of interest (VOIs) were acquired with an isotropic voxel size of 2.21 µm, following the established protocols of Walsh et al. (24). Raw projections were reconstructed into 16-bit TIFF stacks using the ESRF Nabu pipeline (30) and NightRail workflow (31). Following flat-field and dark correction, single-distance Paganin phase retrieval (32) and a Gaussian unsharp mask were applied in preparation for filtered back-projection and ring artifact removal (33).

### Contrast-to-noise ratio quantification

The contrast-to-noise ratio (CNR) was quantified in overview HiP-CT images using an in-house Python script according to:

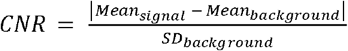

Five independent measurements were obtained from the spinous process, vertebral trabecular bone, paraspinal muscle, and spinal cord using surrounding formalin-agar as the background reference. For comparison, the CNR was calculated from T2w MR images using matched regions of interest and corresponding anatomical locations, with the image background as the reference signal. CNR could not be quantified in the local HiP-CT zoom images as no suitable background region was present within the FOV.

### Segmentation and visualization

Arterial, venous, and neural structures were semi-automatically segmented in overview HiP-CT volumes using a bounded region-growing algorithm in VGSTUDIO MAX (version 2025.3; Volume Graphics, Germany). Vertebral bone was segmented using a global intensity threshold with manual refinement, while IVDs were manually segmented in Avizo3D (version 2022.2; Thermo Fisher Scientific, USA) with interpolation between annotations. Segmentations were volume rendered in VGSTUDIO MAX and Cinematic Anatomy (Siemens Healthineers, Germany).

### IVD grading

IVD degeneration was assessed in midsagittal T2w MR images according to the Miyazaki classification (34) for cervical discs and Pfirrmann classification for thoracic and lumbar discs (35). Two blinded observers (CL and AKGP) performed disc grading independently.

### Fiber tracing and orientation analysis

Annulus fibrosus fiber architecture of the L2-L3 and L4-L5 IVDs was quantified in overview HiP-CT images using Avizo3D (version 2023.2; XFiber module) as previously described (36). Briefly, cylinder correlation and trace correlation lines were used to compute local orientation fields and fiber centerline trajectories within segmented annulus fibrosus volumes, respectively. Fiber orientation was quantified using a custom Python code (https://github.com/UCL-MXI-Bio/Intervertebral-Disc-Fibre-Analysis) relative to the craniocaudal axis and local circumferential direction.

### Vertebral trabecular bone morphometry

Trabecular morphometry was evaluated in cervical (C2-C7) and lumbar (L1-L5) vertebral bodies using the overview HiP-CT dataset. C1 (atlas) was excluded due to the lack of a true vertebral body. Image stacks of each vertebra were anatomically realigned to ensure consistent orientation in DataViewer (version 1.5.6.2; Bruker, Belgium). The trabecular volume was then defined from the first slice immediately below the cranial endplate to the final slice before the caudal endplate and manually segmented in CTAn (version 1.23.0.2+; Bruker, Belgium). Morphometric parameters were subsequently quantified in three dimensions using a custom processing task list in CTAn, with segmented volumes rendered in Avizo3D (version 2023.2).

## Results

### MRI assessment of spinal anatomy and IVD degeneration

The intact spine (C1-L5) was obtained from a 62-year-old male donor who died of stomach cancer and had no known skeletal disorders (Table 1). To provide a conventional anatomical reference for the subsequent HiP-CT assessment, the specimen was first imaged *ex vivo* with MRI using T1w, T2w, multi-echo T2*w and T2w-STIR sequences (Figure 2A). T1w and T2*w MRI delineated gross vertebral and adjacent soft tissue anatomy including prevertebral fat, whereas T2w and STIR imaging enabled fluid-sensitive assessment in vertebral bodies, endplates, IVDs, nucleus pulposus, spinal cord, and cerebrospinal fluid (Figure 2A and B). Midsagittal T2w-MRI-based grading revealed regional variation in IVD degeneration (grades I-IV), with the highest grades observed at C4-C5 and C5-C6, while all lumbar IVDs were assigned a grade of II (Figure 2C).

**Figure 2.**
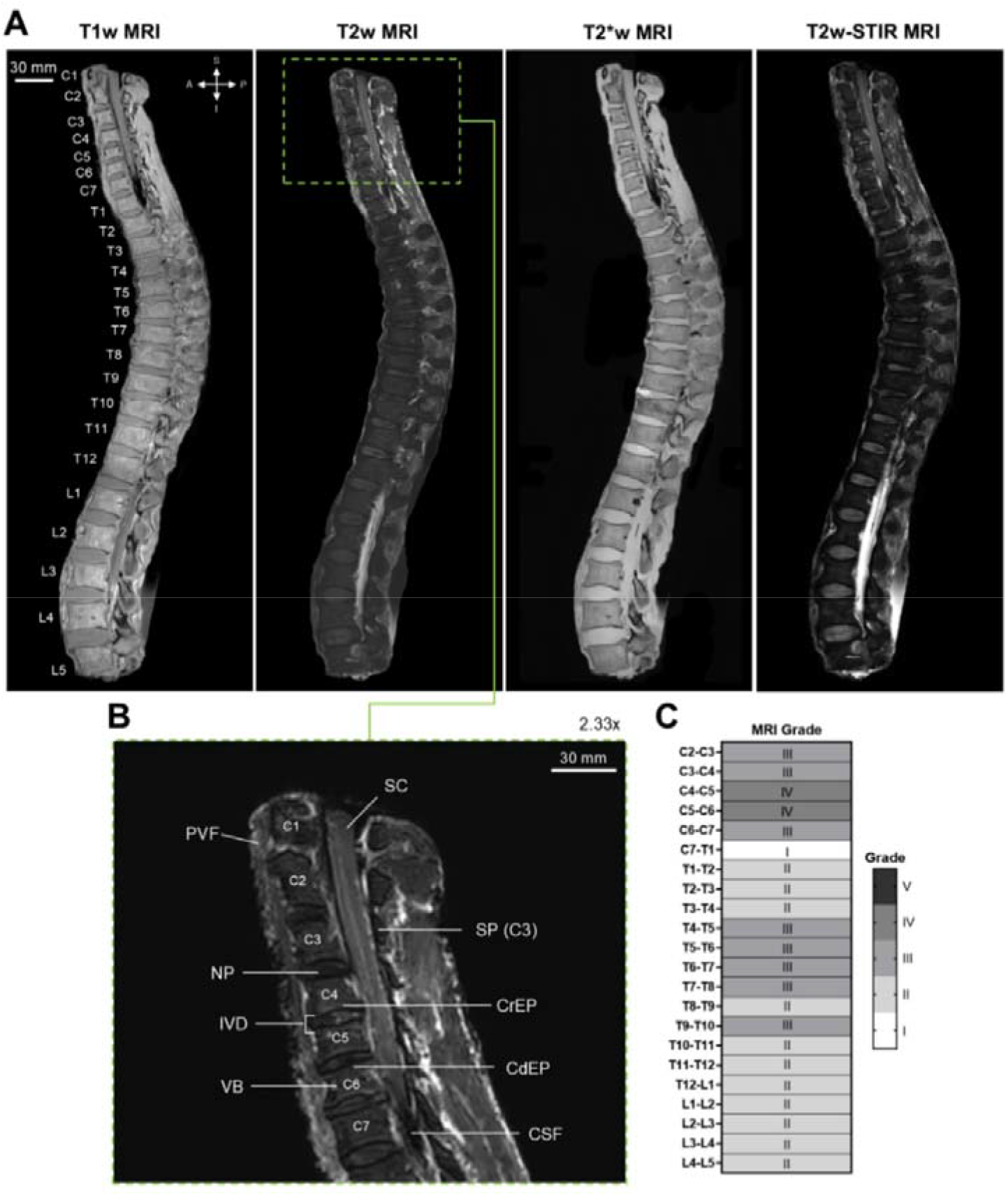
Whole-spine MRI defines anatomy and disc degeneration for HiP-CT region selection. Sagittal 2D T1-weighted (T1w), T2-weighted (T2w), T2*-weighted (T2*w) and T2w-short tau inversion recovery (T2w-STIR) MR images of the intact spine (A). Vertebral levels from C1 to L5 are labeled on the T1w MR image for anatomical reference. Zoomed T2w MR image of the cervical spine highlights vertebral bodies (VB), cranial (CrEP) and caudal endplates (CdEP), intervertebral discs (IVD), nucleus pulposus (NP), spinous processes (SP), spinal cord (SC), cerebrospinal fluid (CSF), and prevertebral fat (PVF, B). MRI-based Miyazaki and Pfirrmann grading showed regional variation in IVD degeneration (C).

### Hierarchical imaging of the whole human spine

HiP-CT enabled high-resolution visualization of the intact spine from C1 to L5 within a single tomographic acquisition, preserving the spatial relationships between mineralized and non-mineralized soft tissues throughout the specimen through endogenous phase-contrast (Figure 3A). At 16.26 µm isotropic voxel size, vertebral cortical and trabecular bone, IVDs, posterior elements, the spinal canal and surrounding soft tissues were simultaneously visualized, with individual trabeculae readily resolved within the vertebral bodies (Figure 3A and B). Local tomography at 2.2 µm isotropic voxel size further resolved Haversian systems within the cranial vertebral endplate (Figure 3C), annulus fibrosus organization (Figure 3D) within the IVDs, as well as facet joint hyaline articular cartilage (Figure 3E) in the lumbar spine. Axial virtual sectioning of the thoracic spine (Figure 3F and G) revealed anatomical relationships not readily appreciated in sagittal MRI, including the segmental vasculature, spinal cord and meninges, ligamentous and posterior element anatomy, multifidus muscle, and the internal organization of the T11-T12 IVD, with the annulus fibrosus and nucleus pulposus clearly visible.

**Figure 3.**
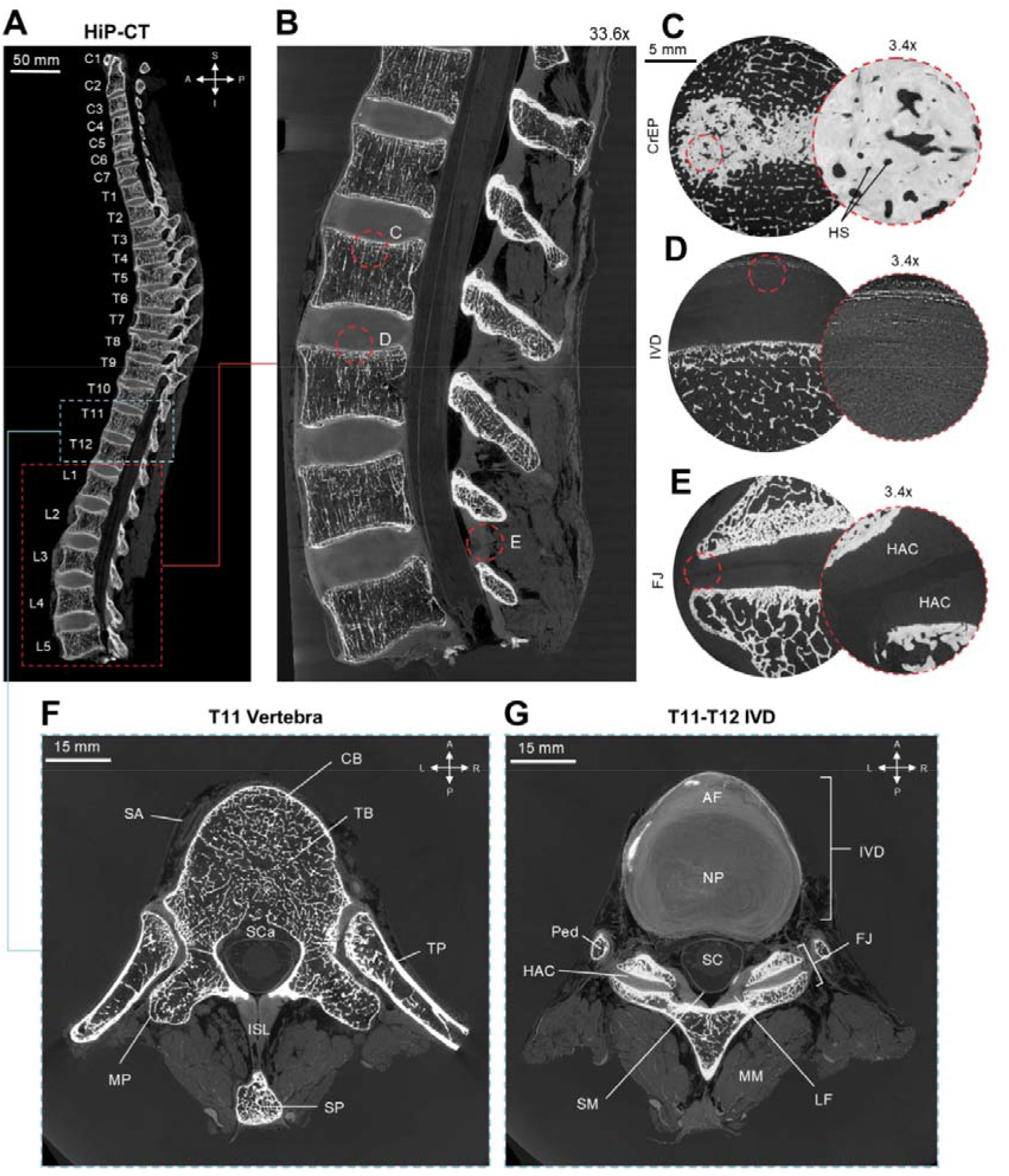
HiP-CT reveals multiscale osseous and soft-tissue microanatomy of the intact human spine. Sagittal overview HiP-CT image of the intact spine acquired using an isotropic voxel size of 16.26 µm (A). Zoomed sagittal view of the lumbar spine (L1-L5) showing vertebral bodies, posterior elements, IVD, and trabecular architecture (B). Representative high-resolution local HiP-CT images, acquired with 2.2 µm isotropic voxels, enable visualization of Haversian systems within the CrEP (C), annulus fibrosus (AF) fiber organization within the IVD (D), and hyaline articular cartilage (HAC) at the facet joint (FJ, E); circular insets show 3.4x digital zooms of regions indicated by the red hatched box in B. Axial HiP-CT overview image of the T11 vertebra (F) showing trabecular bone within the VB and surrounding cortical bone (CB), segmental artery (SA), spinal canal (SCa), transverse process (TP), mammillary process (MP), spinous process (SP), and interspinous ligament (ISL). In the corresponding axial HiP-CT overview image of the T11-T12 IVD, the AF and NP, spinal cord (SC), spinal meninges (SM), HAC within the FJ, pedicle (ped), ligamentum flavum (LF) and multifidus muscle (MF) are also resolved.

Despite the substantially higher spatial resolution of HiP-CT, *ex vivo* MRI demonstrated higher descriptive CNR across all evaluated tissues (Table 2). These modalities therefore showed complementary strengths; MRI provided greater tissue-to-background contrast in this *ex vivo* specimen, whereas HiP-CT resolved micron-scale detail across mineralized and non-mineralized tissues. These CNR differences should be interpreted in the context of *ex vivo* MRI acquisition and the markedly different spatial resolutions of the two modalities.

**Table 2:**
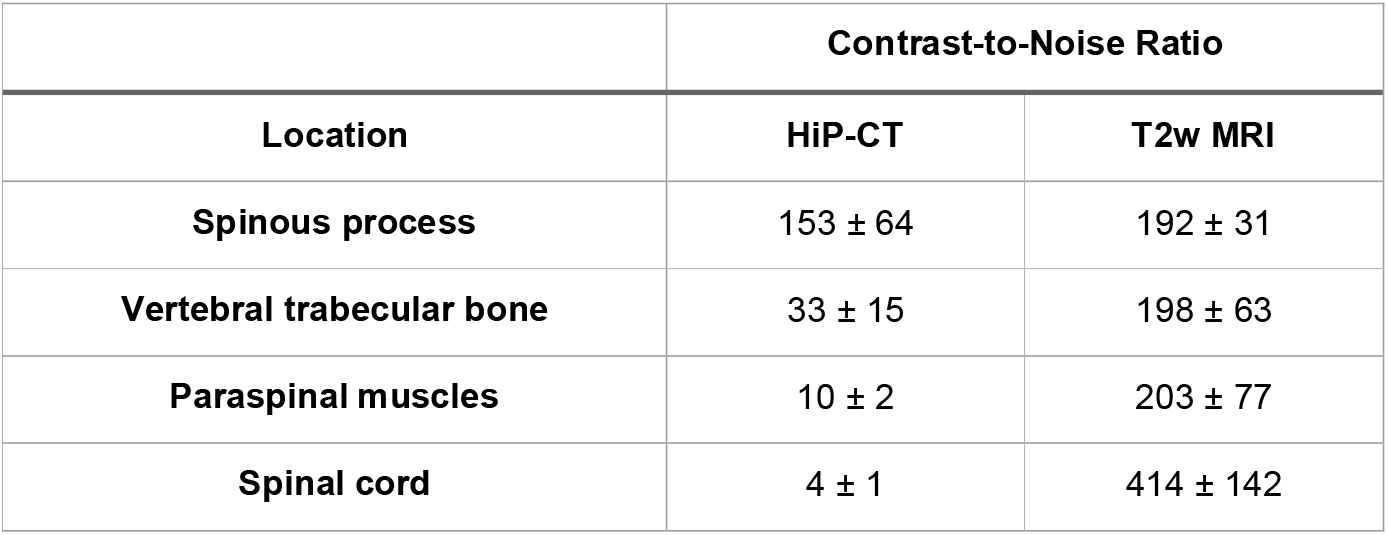
Quantification of the contrast-to-noise (CNR) ratio across selected spinal tissues in HiP-CT and MRI. Values are presented as mean ± SD from five independent measures per tissue.

| Location | Contrast-to-Noise Ratio |  |
| --- | --- | --- |
|  | HiP-CT | T2w MRI |
| Spinous process | 153 ± 64 | 192 ± 31 |
| Vertebral trabecular bone | 33 ± 15 | 198 ± 63 |
| Paraspinal muscles | 10 ± 2 | 203 ± 77 |
| Spinal cord | 4 ± 1 | 414 ± 142 |

### HiP-CT enables the three-dimensional delineation of spinal neurovascular anatomy

HiP-CT provided sufficient endogenous soft tissue contrast to delineate neural and vascular structures in overview images of the intact mineralized vertebral column without exogenous contrast enhancement, enabling their segmentation alongside vertebral bone and IVDs (Figure 4A, Movie S1). Virtual sectioning of these segmented volumes demonstrated their preserved native anatomical relationships, including branching cervical nerve roots, the arterial supply to the thoracic spinal cord, and the vascularization of lumbar vertebral bodies (Figure 4B). By comparison, these finer neurovascular features were not separately discernible on the corresponding *ex vivo* MRI.

**Figure 4:**
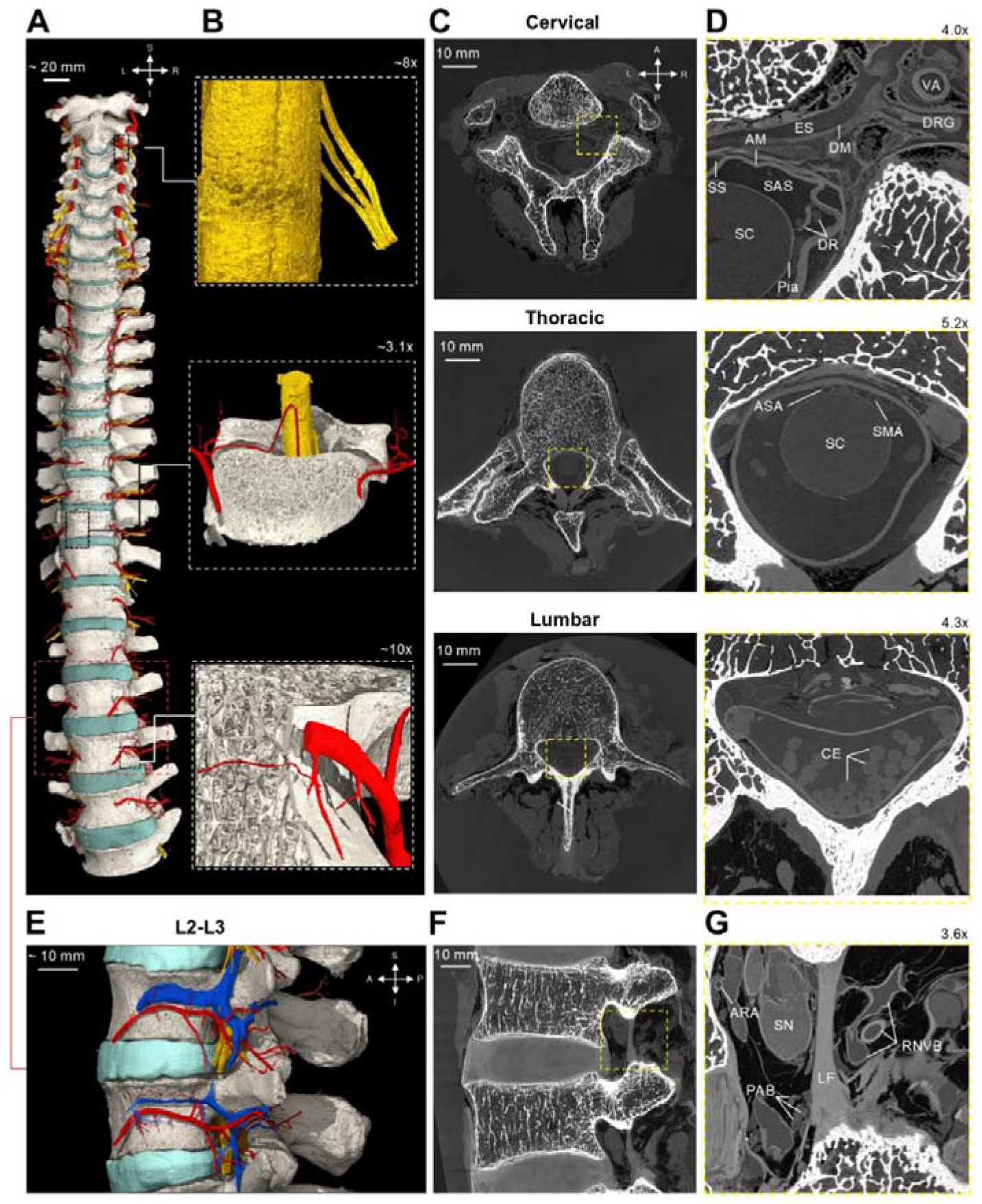
HiP-CT resolves neural and vascular anatomy throughout the intact spine without exogenous contrast agents. Three-dimensional rendering shows segmented vertebrae (white), IVDs (teal), arteries (red), and spinal cord and neural tissues (yellow) along the length of the spine in the coronal plane (A). Virtually sectioned views show branching cervical nerve roots (B, top), thoracic spinal cord vasculature (middle), and vascularization of the L3 vertebral body (bottom). Representative axial HiP-CT overview images of the cervical, thoracic, and lumbar spine (C). Corresponding digital zooms reveal the spinal cord (SC) and surrounding pia mater (Pia), arachnoid membrane (AM), dura mater (DM), epidural space (ES), subarachnoid space (SAS), and subdural space (SS), together with dorsal nerve (DR) rootlets converging towards the dorsal root ganglion (DRG) and the adjacent vertebral artery (VA) in the cervical spine (D, top). The anterior spinal artery (ASA) and segmental medullary artery (SMA) are visible in relation to the thoracic spinal cord (SC; D, middle), while individual cauda equina (CE) nerve roots are resolved within the lumbar spinal canal (D, bottom). Rendering of the lumbar L2-L3 segment shows the three-dimensional distribution of arteries, veins and neural structures relative to the vertebral column and IVD (E), with the corresponding sagittal HiP-CT overview image shown in F. Digital magnification of the lumbar intervertebral foramen (G) reveals the spinal nerve (SN), radicular neurovascular bundle (RNVB), anterior radicular artery (ARA), prelaminar arterial branches (PAB), and ligamentum flavum (LF). Hatched boxes indicate regions shown at higher digital magnification.

Across spinal levels, HiP-CT revealed distinct neurovascular relationships within the 16.26 µm overview images. In the cervical spine, the spinal cord and meningeal layers were visualized in relation to the dorsal nerve rootlets, dorsal root ganglion and adjacent vertebral artery (Figure 4C and D, top). The pia mater was seen to follow the spinal cord surface, with the subarachnoid space extending between the pia and arachnoid membrane, while the dura mater formed the outer meningeal boundary adjacent to the epidural space. In the thoracic spine, intrinsic tissue contrast enabled direct visualization of the anterior spinal and segmental medullary arteries in relation to the spinal cord (Figure 4C and D, middle), while individual cauda equina nerve roots were distinguishable within the lumbar spinal canal (Figure 4C and D, bottom).

At L2-L3, three-dimensional segmentation further demonstrated the spatial organization of arterial, venous, and neural structures relative to the vertebral bodies and IVD (Figure 4E), with corresponding structures directly visible within sagittal HiP-CT overview images (Figure 4F). Within the lumbar intervertebral foramen, the spinal nerve, radicular neurovascular bundle, anterior radicular artery, and prelaminar arterial branches were resolved in relation to the ligamentum flavum and the surrounding osseous boundaries of the foramen (Figure 4G). Venous vasculature was also segmented, although post-mortem vessel collapse limited continuous segmentation beyond this region of the lumbar spine.

### Three-dimensional fiber mapping reveals focal disruption of annulus fibrosus lamellar organization

The overview HiP-CT dataset provided sufficient spatial resolution to trace annulus fibrosus fibers within the lumbar IVDs (Figure 5A and B). The L2-L3 IVD showed well-preserved lamellar organization dominated by circumferentially aligned annulus fibrosus fibers (Figure 5C, pink), whereas the L4-L5 IVD showed bilateral focal posterolateral disruption, with reduced circumferential and increased radial fiber alignment (Figure 5D, yellow). Further quantification of fiber orientation relative to the craniocaudal axis demonstrated a concentration of fibers around ±90° in the L2-L3 IVD (Figure 5E and G). In contrast, the L4-L5 IVD was comprised of fibers that were predominantly angled at approximately ±70° (Figure 5F and H), consistent with a broader and less uniform oblique fiber organization.

**Figure 5.**
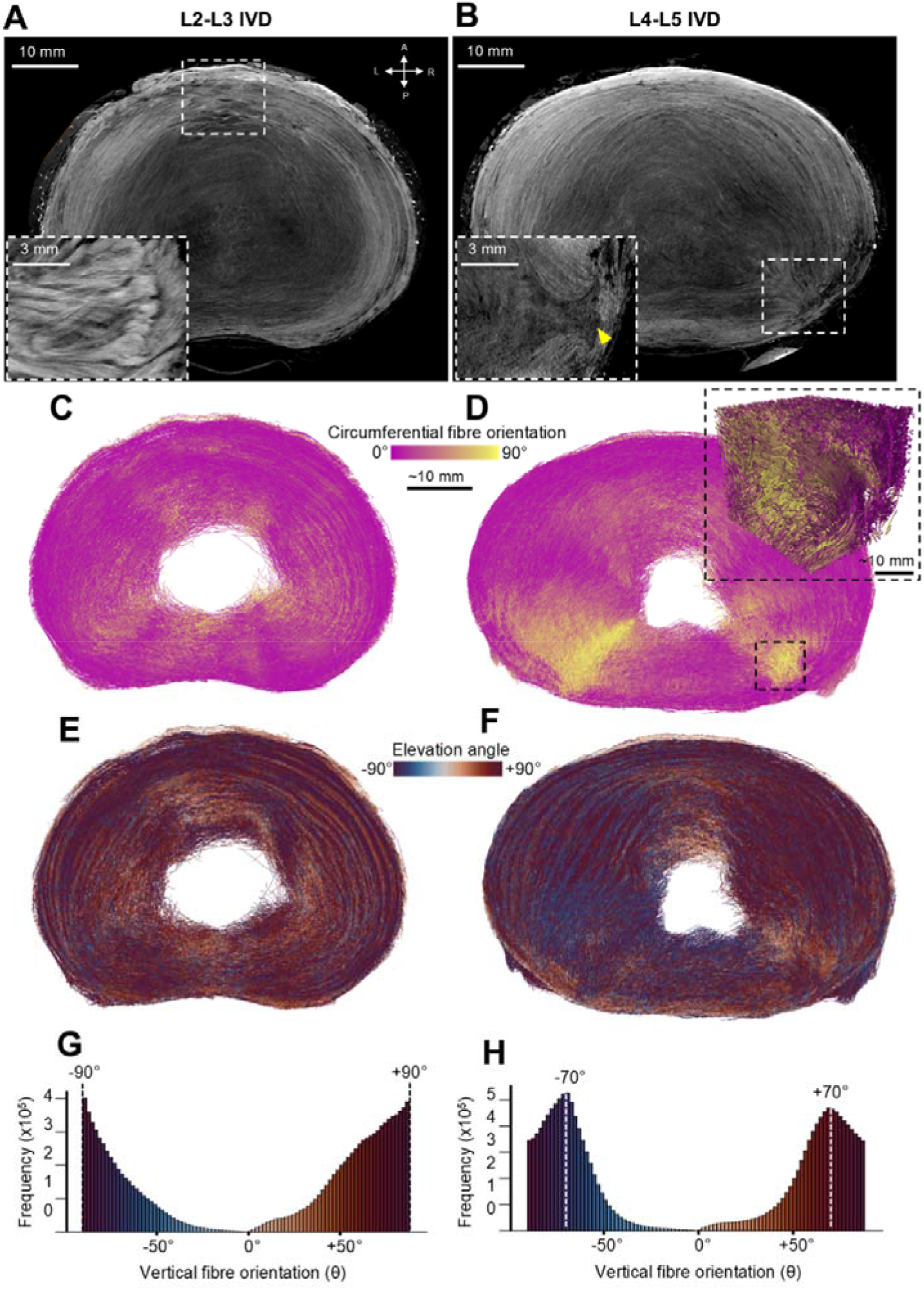
HiP-CT demonstrates regional differences in annulus fibrosus fiber organization between the L2–L3 and L4–L5 intervertebral discs. Mid-axial HiP-CT images from the overview scans of the L2-L3 (A) and L4-L5 (B) IVDs show relatively preserved annulus fibrosus lamellar organization at L2–L3 and focal disruption of the lamellar architecture in the posterolateral region of the L4–L5 IVD (yellow arrowhead). Three-dimensional mapping of fiber orientation relative to the local circumferential direction in the L2–L3 IVD (C) reveals predominant circumferential alignment, whereas increased radial orientation is evident within the focal posterolateral annular disruption in the L4–L5 IVD (D); circumferentially oriented fibers are shown in pink (0°) and radially oriented fibers in yellow (90°). Zoomed volume rendering of annulus fibers within the region of focal disruption in the L4-L5 IVD is shown in the black hatched box. Vertical fiber orientation relative to the craniocaudal axis is shown for L2–L3 (E) and L4–L5 IVDs (F), with corresponding frequency distributions in G and H, respectively. Blue and dark red represent fibers oriented toward –90° and +90°, respectively, highlighting the dominant oblique fiber orientations within each disc.

### HiP-CT permits trabecular morphometric assessment across the cervical and lumbar spine

Three-dimensional HiP-CT renderings demonstrated variation in vertebral body trabecular microarchitecture across cervical and lumbar levels, which were quantified across C2-C7 and L1-L5 (Figure 6A and B). Three-dimensional morphometry revealed that the bone volume fraction (bone volume/tissue volume; BV/TV) was highest at C4-C6, exceeding that at all other cervical and lumbar levels (Figure 6C). Trabecular thickness was comparable between regions (Figure 6D), whereas trabecular spacing was consistently greater in the lumbar than the cervical vertebral bodies. Trabecular number and connectivity density similarly peaked at C4-C6 and were generally greater in cervical than lumbar vertebral bodies (Figure 6F and G). In contrast, trabecular anisotropy was higher in the lumbar vertebral bodies, indicating a more directionally aligned trabecular network, whereas cervical trabeculae were comparatively isotropic (Figure 6H). Additional morphometric parameters are presented in the Supplementary Materials (Figure S2).

**Figure 6.**
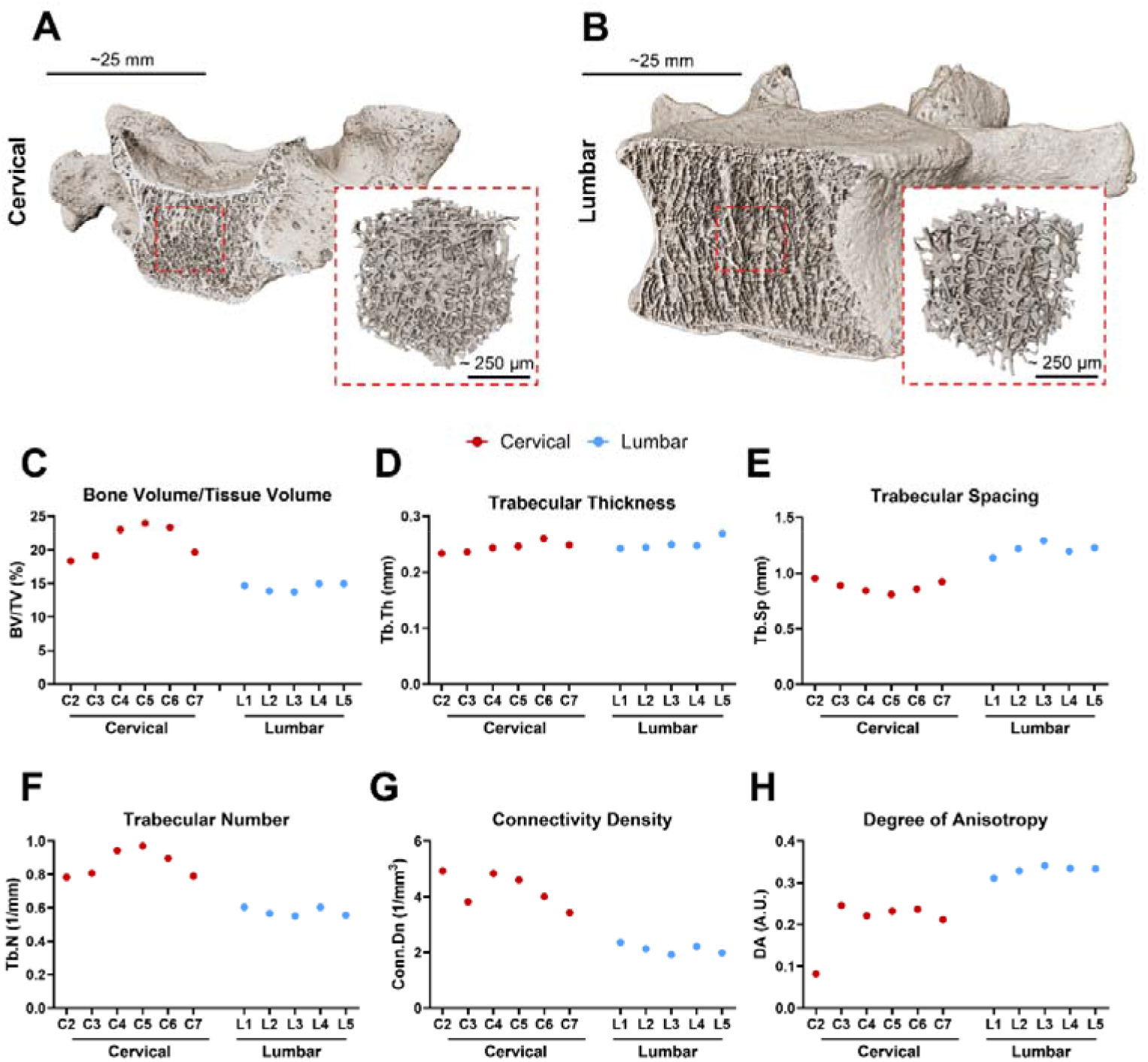
Regional variation in vertebral trabecular microarchitecture across the cervical and lumbar spine. Representative three-dimensional volume rendering of the cervical (C4, A) and lumbar (L3, B) vertebrae with zooms corresponding to trabecular bone within the vertebral bodies generated from the overview HiP-CT scans. Vertebral-level three-dimensional morphometric quantification of trabecular bone volume/tissue volume (C), trabecular thickness (D), trabecular spacing (E), trabecular number (F), connectivity density (G), and degree of anisotropy (H). Across this specimen, the cervical vertebral bodies showed greater trabecular bone volume/tissue volume, trabecular number, and connectivity density whereas lumbar vertebral trabecular bone showed greater trabecular spacing and anisotropy. Trabecular thickness was broadly comparable between regions.

## Discussion

This proof-of-concept study demonstrates that HiP-CT can non-destructively image a whole intact adult human spine *ex vivo,* spanning macro- to microscopic scales at high spatial resolution. Imaging using 16.26 µm isotropic voxels permitted simultaneous visualization of mineralized and non-mineralized tissues, preserving three-dimensional relationships and allowing quantitative analysis of annulus fibrosus fiber organization and vertebral trabecular morphometry. Local tomography using 2.21 µm voxels further resolved annulus fibrosus lamellae, vertebral endplate Haversian systems, and facet joint hyaline cartilage. These findings demonstrate that HiP-CT resolves spinal structure across scales, highlighting its applicability in correlative studies relating features recognizable in clinical imaging to their underlying microarchitectures.

Previous synchrotron X-ray phase-contrast tomography of human spinal specimens achieved 46 µm and 8 µm isotropic resolution; however, visualization of vasculature required perfusion with iodinated contrast agents, while spinal cord microvasculature and gray-white matter architecture could only be assessed following cord dissection (23). Building on this work, HiP-CT extends phase-contrast imaging for the first time to an intact adult human spine at substantially higher resolution, allowing osseous, discal, neural and vascular structures to be visualized together *in situ* without exogenous contrast or tissue dissection. These advances, coupled with the capacity to image broad anatomical regions while preserving native tissue relationships, enable HiP-CT to interrogate microstructural features within their wider anatomical context.

Three-dimensional segmentation demonstrated the feasibility of assessing spinal neurovascular anatomy within the intact vertebral column. Previous HiP-CT studies quantified renal microvascular architecture using vessel caliber, branching geometry, tortuosity and inter-vessel spacing (37), and brain neural architecture using structure tensor and tractography-based approaches (38). Applied to the spine, analogous methods could characterize vascular branching, neural trajectory and organization in relation to the spinal canal and intervertebral foramina. Such analyses could support investigations into vascular compromise, canal or foraminal narrowing and neural compression with potential relevance to surgical or image-guided radiologic assessments.

Beyond anatomical visualization, HiP-CT enabled quantitative assessment of IVD and vertebral trabecular microarchitecture within the same whole-spine dataset. For IVDs, HiP-CT resolved annulus fibrosus organization at spatial scales inaccessible to clinical MRI and enabled fiber tracing throughout intact IVDs; the limited spatial resolution of MRI necessitates reliance on planar assessment of disc signal, morphology, and height (34, 35). In this specimen, relatively preserved circumferential organization of the L2-L3 IVD contrasted with the L4-L5 IVD, where posterolateral disruption was accompanied by increased radial alignment. Notably, all lumbar IVDs were assigned to an MRI-based degenerative grade of II despite this local microarchitectural heterogeneity, suggesting that HiP-CT-informed calibration could refine the relationship between conventional MRI grading and underlying disc structure. Dominant fiber orientations in both discs also extended beyond the previously reported range of ±46°–60° (39), suggesting greater spatial heterogeneity than previously appreciated. Complementing this, HiP-CT also enabled vertebral trabecular morphometric evaluation across multiple spinal levels, highlighting its capacity for integrated visualization and analysis of mineralized and non-mineralized tissues within the same whole-spine dataset. Importantly, this was achieved without specimen dissection typically required in microCT studies (40, 41). Bone volume fraction, trabecular thickness, spacing and structural organization are established determinants of human vertebral mechanical behavior (42), yet this microarchitecture is not directly resolved at comparable spatial scale by clinical imaging. Although differences were observed between cervical and lumbar vertebral bodies, these should be interpreted in the context of this single-donor study. Future studies should examine larger, phenotypically characterized cohorts spanning spinal phenotypes and pathologies, including disc degeneration and vertebral fragility, across IVD grades, using HiP-CT to define tissue-specific microanatomical changes in relation to clinical imaging and histopathological findings.

This study has several limitations. HiP-CT requires synchrotron facilities and technical expertise, restricting throughput and current application to *ex vivo* research as radiation exposure precludes *in vivo* imaging. Multi-terabyte datasets impose substantial storage, visualization, and analytical demands. The single-donor design limits generalizability and precludes statistical analyses. Postmortem fixation and imaging may have influenced tissue hydration, disc morphology, fiber orientation, neurovascular architecture, and MR signal characteristics. Histopathologic validation was unavailable; therefore, the focal posterolateral disruption identified in the L4-L5 IVD should be regarded as a structural observation rather than definitive evidence of disc herniation. Manual and semi-automated image processing may introduce observer bias, and near-micron local tomography was confined to selected regions of interest.

In conclusion, we establish the feasibility of HiP-CT for high-resolution three-dimensional imaging of an intact adult human spine. Whole-spine imaging at 16.26 µm isotropic voxel size, combined with targeted local tomography at 2.21 µm, enables interrogation of spinal macro- to microarchitecture without tissue sectioning or exogenous contrast agents. By integrating anatomical visualization with quantitative analysis across interconnected tissue types, HiP-CT enables *ex vivo* investigations of spinal structure in health and disease. The resulting three-dimensional datasets could constitute a microanatomical reference for relating clinical imaging findings to underlying tissue architecture, supporting the discovery and validation of imaging biomarkers.

## Abbreviations

HiP-CT: Hierarchical phase-contrast tomography
CNR: Contrast-to-noise ratio
IVD: Intervertebral disc
FOV: Field of view
VOI: Volume of interest

## Author contributions

Conceptualization: AS, JB, PT, CLW, PDL; Methodology: AS, ICH, ALP, JB, JC, JP, TU, HD, AD, LL, TJPB, HJW, AB, PT, PDL; Software: AS, ALP, JB, DS, YZ, TU, HD, PT; Validation: AS, ALP, JB; Formal analysis: AS, ICH, ALP, JB, CL, AKGP, JL; Investigation: AS, ICH, JB, JC, JP, TU, HD, AD, LL, AB, PT, PDL; Resources: AB, PDL; Data Curation: AS, JB, DS, YZ, AB, CLW; Writing – Original Draft: AS, ICH, ALP, JB; Writing – Review and editing – All authors; Visualization: AS, ICH, ALP, JB, TU, AAP, TJPB, PDL; Supervision: PDL; Project administration: AS, JB, CLW, PDL; Funding acquisition: PDL.

## Acknowledgements

We acknowledge the ESRF and the HOAHub for provision of synchrotron radiation facilities under UCL-led proposals MD-1290 and MD-1389. This project was made possible in part by grant number 2022-316777 from the Chan Zuckerberg Initiative, MR/V033506/1 from the UK Medical Research Council, and CiET18/19-10 from the Royal Academy of Engineering. Part of this work was performed on the IRMaGe platform member of the France Life Imaging Network under grant number ANR-11-INBS-0006. We gratefully acknowledge the donor and their family whose contribution made this research possible. We also thank Klaus Engel (Siemens Healthineers, Erlangen, Germany) for his assistance in generating the cinematic anatomical renderings used in this study.

## Data availability

The HiP-CT imaging datasets generated in this study will be deposited in the HOAHub repository (https://human-organ-atlas.esrf.fr/) and made publicly available upon publication of the peer-reviewed article.

## References

1. Ghodasara N, Yi PH, Clark K, Fishman EK, Farshad M, Fritz J. Postoperative Spinal CT: What the Radiologist Needs to Know. Radiographics 2019;39(6):1840–1861. doi: 10.1148/rg.2019190050

2. Van Goethem JW, Maes M, Ozsarlak O, van den Hauwe L, Parizel PM. Imaging in spinal trauma. Eur Radiol 2005;15(3):582–590. doi: 10.1007/s00330-004-2625-5

3. Kalichman L, Kim DH, Li L, Guermazi A, Hunter DJ. Computed tomography-evaluated features of spinal degeneration: prevalence, intercorrelation, and association with self-reported low back pain. Spine J 2010;10(3):200–208. doi: 10.1016/j.spinee.2009.10.018

4. Isikbay M, Talbott J. Clinical Guidelines and Their Role in Trauma-related Cervical Spine Imaging. Radiology 2026;318(1):e252327. doi: 10.1148/radiol.252327

5. Winegar BA, Kay MD, Taljanovic M. Magnetic resonance imaging of the spine. Pol J Radiol 2020;85:e550–e574. doi: 10.5114/pjr.2020.99887

6. Kim GU, Chang MC, Kim TU, Lee GW. Diagnostic Modality in Spine Disease: A Review. Asian Spine J 2020;14(6):910–920. doi: 10.31616/asj.2020.0593

7. Sollmann N, Fields AJ, O’Neill C, Nardo L, Majumdar S, Chin CT, Tosun D, Han M, Vu AT, Ozhinsky E, Shah LM, Harris RE, Lobo R, Anderst W, Herzog R, Psioda MA, Standaert CJ, Price RT, Lotz JC, Link TM, Krug R. Magnetic Resonance Imaging of the Lumbar Spine: Recommendations for Acquisition and Image Evaluation from the BACPAC Spine Imaging Working Group. Pain Med 2022;24(Supplement_1):S81–S94. doi: 10.1093/pm/pnac130

8. Deininger-Czermak E, Villefort C, von Knebel Doeberitz N, Franckenberg S, Kälin P, Kenkel D, Gascho D, Piccirelli M, Finkenstaedt T, Thali MJ, Guggenberger R. Comparison of MR Ultrashort Echo Time and Optimized 3D-Multiecho In-Phase Sequence to Computed Tomography for Assessment of the Osseous Craniocervical Junction. J Magn Reson Imaging 2021;53(4):1029–1039. doi: 10.1002/jmri.27478

9. Schwaiger BJ, Schneider C, Kronthaler S, Gassert FT, Böhm C, Pfeiffer D, Baum T, Kirschke JS, Karampinos DC, Makowski MR, Woertler K, Wurm M, Gersing AS. CT-like images based on T1 spoiled gradient-echo and ultra-short echo time MRI sequences for the assessment of vertebral fractures and degenerative bone changes of the spine. Eur Radiol 2021;31(7):4680–4689. doi: 10.1007/s00330-020-07597-9

10. Rau A, Straehle J, Stein T, Diallo T, Rau S, Faby S, Nikolaou K, Schoenberg SO, Overhoff D, Beck J, Urbach H, Klingler JH, Bamberg F, Weiss J. Photon-Counting Computed Tomography (PC-CT) of the spine: impact on diagnostic confidence and radiation dose. Eur Radiol 2023;33(8):5578–5586. doi: 10.1007/s00330-023-09511-5

11. Mahler L, Steiglechner J, Bender B, Lindig T, Ramadan D, Bause J, Birk F, Heule R, Charyasz E, Erb M, Kumar VJ, Hagberg GE, Martin P, Lohmann G, Scheffler K. Submillimeter Ultra-High Field 9.4 T Brain MR Image Collection and Manual Cortical Segmentations. Sci Data 2025;12(1):635. doi: 10.1038/s41597-025-04779-2

12. Wu LL, Huang SF, Zhu LH, Liu H, Zhou JJ. Quantitative evaluation of lumbar intervertebral disc degeneration: a comparison of ultrashort time-to-echo T2* with T1rho relaxometry. Acta Radiol 2025;66(2):137–145. doi: 10.1177/02841851241309234

13. Gassert FT, Kufner A, Gassert FG, Leonhardt Y, Kronthaler S, Schwaiger BJ, Boehm C, Makowski MR, Kirschke JS, Baum T, Karampinos DC, Gersing AS. MR-based proton density fat fraction (PDFF) of the vertebral bone marrow differentiates between patients with and without osteoporotic vertebral fractures. Osteoporos Int 2022;33(2):487–496. doi: 10.1007/s00198-021-06147-3

14. Ellingson BM, Ulmer JL, Kurpad SN, Schmit BD. Diffusion Tensor MR Imaging of the Neurologically Intact Human Spinal Cord. AJNR Am J Neuroradiol 2008;29(7):1279–1284. doi: 10.3174/ajnr.A1064

15. Guo Y, Liu Z, Wen Y, Spincemaille P, Zhang H, Jafari R, Zhang S, Eskreis-Winkler S, Gillen KM, Yi P, Feng Q, Feng Y, Wang Y. Quantitative susceptibility mapping of the spine using in-phase echoes to initialize inhomogeneous field and R2* for the nonconvex optimization problem of fat-water separation. NMR Biomed 2019;32(11):e4156. doi: 10.1002/nbm.4156

16. Irie TY, Irie T, Espinoza Orías AA, Segami K, Iwasaki N, An HS, Inoue N. Micro-computed tomography analysis of the lumbar pedicle wall. PLoS One 2021;16(7):e0253019. doi: 10.1371/journal.pone.0253019

17. Buckles MM, Nur A, Warda NH, Moncher JC, Ansari MY. High-Resolution 3-Dimensional Micro-CT Imaging of Intervertebral Discs Using a Novel Contrast Agent. JOR SPINE 2025;8(4):e70125. doi: 10.1002/jsp2.70125

18. de Bournonville S, Vangrunderbeeck S, Kerckhofs G. Contrast-Enhanced MicroCT for Virtual 3D Anatomical Pathology of Biological Tissues: A Literature Review. Contrast Media Mol Imaging 2019;2019:8617406. doi: 10.1155/2019/8617406

19. Goyal D, Kumar H. In Vivo and 3D Imaging Technique(s) for Spatiotemporal Mapping of Pathological Events in Experimental Model(s) of Spinal Cord Injury. ACS Chem Neurosci 2023;14(5):809–819. doi: 10.1021/acschemneuro.2c00643

20. Kellner M, Heidrich M, Lorbeer R-A, Antonopoulos GC, Knudsen L, Wrede C, Izykowski N, Grothausmann R, Jonigk D, Ochs M, Ripken T, Kühnel MP, Meyer H. A combined method for correlative 3D imaging of biological samples from macro to nano scale. Sci Rep 2016;6(1):35606. doi: 10.1038/srep35606

21. Cao Y, Wu T, Yuan Z, Li D, Ni S, Hu J, Lu H. Three-dimensional imaging of microvasculature in the rat spinal cord following injury. Sci Rep 2015;5:12643. doi: 10.1038/srep12643

22. Yeh S-CA, Wilk K, Lin CP, Intini G. In Vivo 3D Histomorphometry Quantifies Bone Apposition and Skeletal Progenitor Cell Differentiation. Sci Rep 2018;8(1):5580. doi: 10.1038/s41598-018-23785-6

23. Barbone GE, Bravin A, Mittone A, Grosu S, Ricke J, Cavaletti G, Djonov V, Coan P. High-Spatial-Resolution Three-dimensional Imaging of Human Spinal Cord and Column Anatomy with Postmortem X-ray Phase-Contrast Micro-CT. Radiology 2021;298(1):135–146. doi: 10.1148/radiol.2020201622

24. Walsh CL, Tafforeau P, Wagner WL, Jafree DJ, Bellier A, Werlein C, Kühnel MP, Boller E, Walker-Samuel S, Robertus JL, Long DA, Jacob J, Marussi S, Brown E, Holroyd N, Jonigk DD, Ackermann M, Lee PD. Imaging intact human organs with local resolution of cellular structures using hierarchical phase-contrast tomography. Nat Methods 2021;18(12):1532–1541. doi: 10.1038/s41592-021-01317-x

25. Vijayakumar J, Dejea H, Mirone A, Muzelle C, Meyer J, Jarnias C, Dollman K, Zabler S, Paolasini L, Bellier A, Cordonnier B, Fernandez V, Tafforeau P. Multiresolution Phase-Contrast Tomography on BM18, a New Beamline at the European Synchrotron Radiation Facility. Synchrotron Radiat News 2024;37(5):16–25. doi: 10.1080/08940886.2024.2414724

26. Brunet J, Cook AC, Walsh CL, Cranley J, Tafforeau P, Engel K, Arthurs O, Berruyer C, Burke O’Leary E, Bellier A, Torii R, Werlein C, Jonigk DD, Ackermann M, Dollman K, Lee PD. Multidimensional Analysis of the Adult Human Heart in Health and Disease Using Hierarchical Phase-Contrast Tomography. Radiology 2024;312(1):e232731. doi: 10.1148/radiol.232731

27. Brunet J, Walsh CL, Wagner WL, Bellier A, Werlein C, Marussi S, Jonigk DD, Verleden SE, Ackermann M, Lee PD, Tafforeau P. Preparation of large biological samples for high-resolution, hierarchical, synchrotron phase-contrast tomography with multimodal imaging compatibility. Nat Protoc 2023;18(5):1441–1461. doi: 10.1038/s41596-023-00804-z

28. Xian RP, Walsh CL, Verleden SE, Wagner WL, Bellier A, Marussi S, Ackermann M, Jonigk DD, Jacob J, Lee PD, Tafforeau P. A multiscale X-ray phase-contrast tomography dataset of a whole human left lung. Sci Data 2022;9(1):264. doi: 10.1038/s41597-022-01353-y

29. Chourrout M, Keenlyside A, Wanjau E, Balbastre Y, Yagis E, Brunet J, Stansby D, Engel K, Gui X, Thönnißen J, Dickscheid T, Lamalle L, Bellier A, Vivekananda U, Tafforeau P, Lee PD, Walsh CL. The Extremely Brilliant Brain: An Isotropic Microscale Human Brain Dataset. bioRxiv 2026. doi: 10.64898/2026.01.27.702076

30. Paleo P, Lesaint J, Payno H, Mirone A, Vigano N, Nemoz C. Nabu 2024.1. Zenodo, 2024. doi: 10.5281/zenodo.11104029

31. Mirone A, Brunet J, Urban T, Dejea H, Admans L, Boistel R, Sowinski M, Paleo P, Payno H, Verleden SE, Berruyer C, Boller E, Walsh CL, Lee PD, Tafforeau P. Eikonal Phase Retrieval: unleashing the potential of fourth-generation sources for enhanced propagation-based tomography on biological samples. J Synchrotron Radiat 2025;32(5):1291–1301. doi: 10.1107/S1600577525005223

32. Paganin D, Mayo SC, Gureyev TE, Miller PR, Wilkins SW. Simultaneous phase and amplitude extraction from a single defocused image of a homogeneous object. J Microsc 2002;206(Pt 1):33–40. doi: 10.1046/j.1365-2818.2002.01010.x

33. Lyckegaard A, Johnson G, Tafforeau P. Correction of ring artifacts in X-ray tomographic images. International Journal of Tomography and Statistics 2011;18(F11):1–9.

34. Miyazaki M, Hong S, Yoon S, Morishita Y, Wang J. Reliability of a Magnetic Resonance Imaging-based Grading System for Cervical Intervertebral Disc Degeneration. J Spinal Disord Tech 2008;21(4):288–292. doi: 10.1097/BSD.0b013e31813c0e59

35. Pfirrmann C, Metzdorf A, Zanetti M, Hodler J, Boos N. Magnetic Resonance Classification of Lumbar Intervertebral Disc Degeneration. Spine 2001;26(17):1873–1878. doi: 10.1097/00007632-200109010-00011

36. Newham E, Parmenter AL, Chen J, Disney C, Snow T, Brunet J, Vinci V, Drnec J, Sherratt MJ, Hoyland JA, Bellier A, Bay BK, Terrill N, Lee PD, Gupta HS. Multimodal X-ray imaging reveals hierarchical fibre mechanics. bioRxiv 2025. doi: 10.1101/2025.09.19.677294

37. Rahmani S, Jafree DJ, Lee PD, Tafforeau P, Brunet J, Nandanwar S, Zhou Y, Jacob J, Bellier A, Ackermann M, Jonigk DD, Shipley RJ, Long DA, Walsh CL. Mapping the arterial vascular network in an intact human kidney using hierarchical phase-contrast tomography. npj Imaging 2025;3(1):39. doi: 10.1038/s44303-025-00090-2

38. Wanjau E, Chourrout M, Maffei C, Balbastre Y, Keenlyside A, Brunet J, Sharma A, Huang SY, Tafforeau P, Fischl B, Yendiki A, Lee PD, Walsh CL. Bridging the microstructural gap in human connectomics using hierarchical phase-contrast tomography as a reference for diffusion MRI in the human brain. bioRxiv 2026. doi: 10.64898/2026.04.02.715729

39. Sharabi M, Wade K, Haj-Ali R. Chapter 7 - The Mechanical Role of Collagen Fibers in the Intervertebral Disc. In: Galbusera F, Wilke H-J, eds. Biomechanics of the Spine: Academic Press, 2018; p. 105–123. doi: 10.1016/B978-0-12-812851-0.00007-0

40. Li K, Yang Y, Wang P, Song H, Ma C, Zhang Y, Dang X, Shi J, Zhang S, Li Z, Wang X. Exploring the micromorphological characteristics of adult lower cervical vertebrae based on micro-computed tomography. Sci Rep 2023;13(1):12400. doi: 10.1038/s41598-023-39703-4

41. Wang Y, Owoc JS, Boyd SK, Videman T, Battié MC. Regional variations in trabecular architecture of the lumbar vertebra: Associations with age, disc degeneration and disc space narrowing. Bone 2013;56(2):249–254. doi: 10.1016/j.bone.2013.06.022

42. Roux JP, Wegrzyn J, Arlot ME, Guyen O, Delmas PD, Chapurlat R, Bouxsein ML. Contribution of Trabecular and Cortical Components to Biomechanical Behavior of Human Vertebrae: An Ex Vivo Study. J Bone Miner Res 2010;25(2):356–361. doi: 10.1359/jbmr.090803

