## Supplementary Material for "Revealing multiscale architecture of the intact human spine using hierarchical phase-contrast tomography"

**Movie legends**

**Movie S1:** Cinematic 3D rendering of the HiP-CT overview scan of the intact human spine from C1 to L5 acquired using an isotropic voxel size of 16.26 µm. Rotating views and virtual clipping planes reveal the vertebral bodies, posterior elements, intervertebral discs, spinal canal and cord with nerve roots (yellow) and associated vasculature (arteries, red; veins, blue) created using Cinematic Anatomy (Siemens Healthineers, Germany).

**Methods**

**MR Image acquisition**

MRI was performed at IRMaGe, Grenoble Central University Hospital. Prior to imaging, the spine was wrapped in plastic film to contain the formalin to prevent leakage during acquisition. Each MRI sequence was acquired using the posterior part of the head-and-neck coil and the table-integrated posterior coil.

**Table S1: MRI image acquisition parameters.** Sagittal 2D T1w, T2w, T2*w and T2w-STIR sequences were acquired using a 3 T Achieva dStream.

|  | T_1_w MRI | T_2_w MRI | T_2_*w MRI | T_2_w-STIR MRI |
| --- | --- | --- | --- | --- |
| Acquisition type | 2D | 2D | 2D | 2D |
| Acquisition plane | Sagittal | Sagittal | Sagittal | Sagittal |
| Sequence type | Spin echo | Spin echo | Gradient echo | Inversion recovery |
| Repetition time, TR (ms) | 529.50 | 3964.34 | 2335.50 | 6890.34 |
| Echo time, TE (ms) | 6.36 | 110 | 2.3, 5.8, 9.2, 12.7, 16.1, 19.6, 23, 26.5, 29.9, 33.4, 36.8, 40.3 | 50 |
| Inversion time, TI (ms) | - | - | - | 120 |
| Flip angle (°) | 90 | 90 | 90 | 90 |
| In plane pixel size (mm) | 0.99 x 1.0 | 0.90 x 1.14 | 0.99 x 1.0 | 0.99 x 1.0 |
| Reconstructed in plane pixel size (mm) | 0.47 x 0.47 | 0.53 × 0.53 | 0.47 x 0.47 | 0.47 x 0.47 |
| Slice thickness (mm) | 2 | 3.5 | 2 | 3.5 |
| Slice separation (mm) | 2 | 3.85 | 2 | 3.5 |
| Field of view (mm) | 226 x 226 | 512 x 512 | 226 x 226 | 226 x 226 |
| Pixel bandwidth (Hz/pixel) | 358 | 250 | 857 | 216 |
| Averages | 1 | 1 | 1 | 1 |
| Number of images in acquisition | 55 | 31 | 67 | 31 |

**Sample mounting for HiP-CT**

Prior to HiP-CT imaging, the intact spine was mounted directly in a polycarbonate tube (80 cm length, 15.9 cm outer diameter and 3 mm wall thickness) and stabilized in formalin-agar. This primary tube was housed within a larger secondary polycarbonate tube (60 cm length, 25.4 cm outer diameter, and 5 mm wall thickness). To minimize radiation-induced bubble formation during imaging, the mounted specimen underwent in-line degassing before the primary polycarbonate tube was hermetically sealed, as described by Walsh et al. (1). Whole-spine overview imaging and local tomography of VOIs 1-4 were acquired with this mounting configuration using a bespoke large-capacity rotation stage (LAB Motion Systems, Belgium) to accommodate the intact specimen. VOIs 6-9 were acquired during a subsequent imaging session. For these acquisitions, the lumbar segment (L1-L5) was removed and remounted in formalin-agar within a cylindrical polyethylene terephthalate jar (25.5 cm in length, 14.4 cm diameter), degassed and sealed as described above, and housed within a polycarbonate tube (56 cm length, 15.9 cm outer diameter, 3 mm wall thickness) for imaging following the protocols as previously described (1, 2).

**HiP-CT image acquisition**

A filtered, parallel polychromatic X-ray beam was used for all HiP-CT acquisitions. The source-to-sample distance was 174 m for whole-spine overview imaging and local tomography of VOIs 1-4, and 178 m for VOIs 6-9. Sample-to-detector propagation distances were 32.3 m for whole-spine overview imaging and 1 m for all local tomographic acquisitions. Further details on acquisition-specific parameters including attenuators, scan time and dose rate are provided in Table S2.

Whole-spine overview images were acquired using continuous helical tomography in half-acquisition fly-scan mode (3), with 15 000 projections collected over a 360° rotation at angular increments of 0.024°, an exposure time of 8 milliseconds per projection, and a z-step of 4.845 mm. The FOV was approximately 150.5 mm in diameter and 649.3 mm in height. Local acquisitions were performed as z-series using the same half-acquisition fly-scan mode as described above, with 15 000 projections collected over a 360° rotation at angular increments of 0.024°, an exposure time of 35 milliseconds per projection, and z-step of 5 mm. The FOV was approximately 19.9-20 mm in diameter and 6.5-21.5 mm in height depending on the VOI.

Transmitted X-rays were converted to visible light using bespoke LuAG:Ce scintillators (Crytur, Czechia) and detected using an Iris 15 sCMOS camera (Teledyne Photometrics, USA). Overview imaging used a 2000 µm scintillator coupled to 0.125x demagnification optics, whereas local tomographic imaging used a 50 µm reflective scintillator coupled to fixed 2x magnification optics. The overview and local acquisition scans resulted in estimated radiation doses of 3.61 kGy and 19-36.9 kGy, respectively.

**HiP-CT image reconstruction and post-processing**

For each acquisition, reference scans of an equivalent container filled with water were acquired prior to and following HiP-CT imaging for flat-field correction and suppression of low-frequency artefacts, with additional dark images acquired in the absence of the X-ray beam for dark correction. Single-distance Paganin-phase retrieval was applied using a δ/β ratio of 1000, followed by a Gaussian unsharp mask (overview scans: σ=1.2 and coefficient=4; local tomography: σ=2 and coefficient=4) and ring artefact removal. Reconstructed volumes were subsequently cropped and converted to 16-bit TIFF image stacks.

**Table S2: HiP-CT acquisition parameters.**

| Context | Voxel size  (µm) | Propagation distance (m) | Mode | Mean energy (keV) | Detector area  (pixels) | Number of projections | Field of view (mm) | Attenuators | Dose rate (Gy/s) | Total scan time  (h) | Dataset size  (TB) |
| --- | --- | --- | --- | --- | --- | --- | --- | --- | --- | --- | --- |
| Whole-spine overview | 16.26 | 32.3 | Half acquisition, helical | 160 | 5056 x 600 | 15000 | 150.5 x 649.3 | 5 mm sapphire,  0.3 mm silver | 23.2 | 4.9 | 6.84 |
| VOI-1  T5 vertebral body | 2.21 | 1 | Half acquisition, zseries | 87 | 5056 x 2960 | 15000 | 19.9 x 16.5 | 2.5 mm sapphire,  0.1 mm silver | 18.0 | 0.5 | 1.23 |
| VOI-2  T6 endplate | 2.21 | 1 | Half acquisition, zseries | 87 | 5056 x 2960 | 15000 | 19.9 x 16.5 | 2.5 mm sapphire, 0.1 mm silver | 18.0 | 0.5 | 1.22 |
| VOI-3  T8 endplate | 2.21 | 1 | Half acquisition, zseries | 87 | 5056 x 2960 | 15000 | 19.9 x 6.53 | 2.5 mm sapphire, 0.1 mm silver | 18.0 | 0.5 | 0.48 |
| VOI-4  T10 endplate-disc junction | 2.21 | 1 | Half acquisition, zseries | 87 | 5056 x 2960 | 15000 | 19.9 x 21.5 | 2.5 mm sapphire, 0.1 mm silver | 18.0 | 0.6 | 1.59 |
| VOI-6  L3 endplate-disc junction | 2.21 | 1 | Half acquisition, zseries | 109.4 | 5056 x 2960 | 15000 | 20.0 x 21.5 | 5 mm sapphire,  0.3 mm silver | 8.9 | 0.6 | 1.60 |
| VOI-7  L5 endplate-disc junction | 2.21 | 1 | Half acquisition, zseries | 109.4 | 5056 x 2960 | 15000 | 20.0 x 16.6 | 5 mm sapphire, 0.3 mm silver | 8.9 | 0.5 | 1.23 |
| VOI-8  L4-5 facet joint | 2.21 | 1 | Half acquisition, zseries | 109.4 | 5056 x 2960 | 15000 | 20.0 x 16.5 | 5 mm sapphire, 0.3 mm silver | 8.9 | 0.5 | 1.23 |
| VOI-9  L2 vertebral body | 2.21 | 1 | Half acquisition, zseries | 109.4 | 5056 x 2960 | 15000 | 20.0 x 16.5 | 5 mm sapphire, 0.3 mm silver | 8.9 | 0.5 | 1.23 |

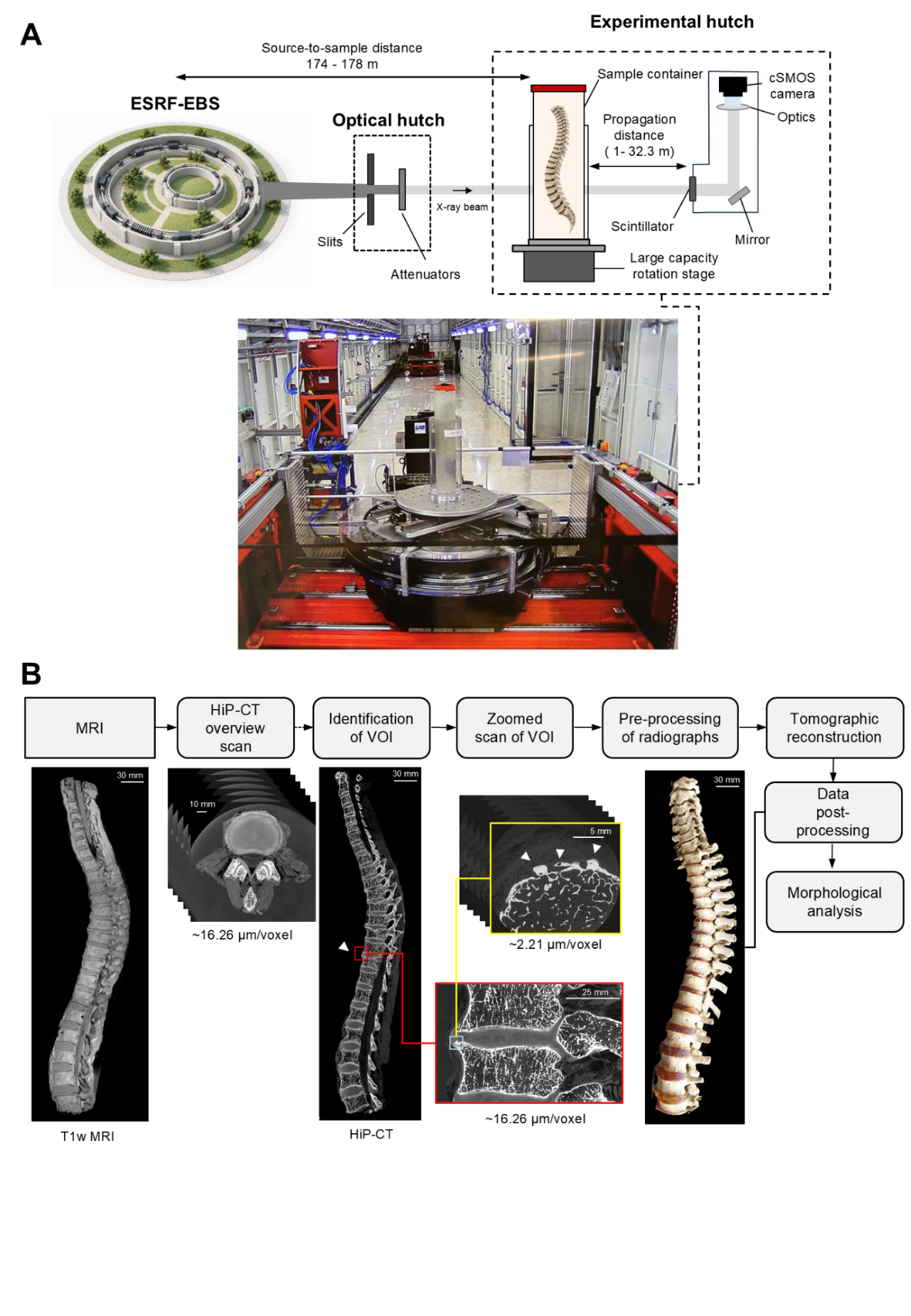

**Figure S1: Ex vivo HiP-CT imaging workflow for the intact human spine**. Schematic of the ESRF BM18 beamline configuration used for HiP-CT imaging, illustrating the source-to-sample distance, optical hutch components (including slits and attenuators) and experimental hutch setup (including scintillators, optics, and detector; A). Schematic of workflow (B) in which MRI images of the intact spine were collected prior to overview HiP-CT scans using isotropic voxels of 16.26 µm with a propagation distance of 32.3 m. Following the identification of VOIs, further high-resolution local HiP-CT scans were performed using 2.2 µm isotropic voxels and propagation distance of 1 m, in combination with alternative optics and a scintillator. All radiographs were flatfield and dark corrected prior to intensity normalization to the synchrotron current ahead of Paganin-phase retrieval and tomographic reconstruction. Reconstructed volumes underwent post-processing to enable hierarchical morphological analysis across the length of the spine.

**Fibre tracing and orientation analysis**

Cylinder correlation was first applied to the image data to generate local correlation and orientation fields for fibre-like structures using a cylinder length of 40 voxels, 5° angular sampling, a mask radius of 3 voxels, outer and inner radii of 2 and 0 voxels, respectively, and bright-on-dark contrast. For the generation of fibre centerline trajectories, the Trace Correlation Lines function was employed and used a minimum seed correlation of 80, minimum continuation quality of 60, direction coefficient of 0.4, minimum fibre spacing of 4 voxels, minimum fibre length of 500, search cone length of 40 voxels, search cone angle of 37°, and minimum step size of 10%.

The annulus fibrosus was manually segmented in 813 µm increments with interpolation between annotations, while interlamellar spaces were excluded by interactive thresholding. The resulting segmentation was used to mask the correlation and orientation fields before tracing, thereby restricting fibre detection to the annulus fibrosus.

For the quantification of fibre orientation, points were seeded at 4 voxel-increments along each traced fibre, and the local fibre tangent was calculated at each point. Fibre elevation (θ) was defined as the angle between the fibre tangent and the craniocaudal axis of the IVD. As the traced fibre centerlines have no inherent directional polarity, 0° represented vertical alignment and ±90° represented horizontal alignment. Fibre orientation relative to the local circumferential direction of each IVD was calculated from the segmented annulus fibrosus geometry. Circumferential reference contours were generated from the inner and outer annulus boundaries on each axial image slice, and the local circumferential direction was defined by the tangent to these contours. Each sampled fibre point was assigned the local circumferential direction of the nearest reference contour, and the angular difference between the fibre tangent and the local circumferential direction was calculated in the axial plane. Circumferential orientation angles therefore ranged from 0° to 90°, with 0° indicating circumferential alignment and 90° indicating radial alignment.

**Vertebral trabecular bone morphometry**

For representative assessment of trabecular bone microarchitecture, cervical (C2-C7) and lumbar (L1-L5) vertebral bodies were selected for the sampling of anatomically distinct spinal regions and to demonstrate the feasibility of quantitative morphometric analysis using the overview HiP-CT images. Vertebral trabeculae were manually segmented in aligned volumes in 244 µm increments using CTAn, prior to binarization using an intensity threshold of 145-255, followed by a 10-pixel black and white despeckling operation to remove isolated foreground and background noise.

**Results**

Additional three-dimensional morphometric analyses demonstrated regional variation in vertebral trabecular microarchitecture (Figure S2). In this specimen, tissue volume (Figure S2A), bone volume (Figure S2B), tissue surface (Figure S2C), and bone surface (Figure S2D) were greater across the lumbar than cervical vertebral bodies consistent with larger anatomical dimensions of L1-L5 vertebrae. Bone surface/tissue volume was consistently greater in the cervical than the lumbar spine (Figure S2E) whereas the bone surface/bone volume (Figure S2F) showed a minor difference between the spinal levels. Trabecular pattern factor and structure model index showed modest level-dependent variation, with lower values across the mid-cervical spine from C4 to C6 (Figure S2G and H, respectively) while fractal dimension remained relatively consistent across the cervical and lumbar vertebral bodies (Figure S2I).

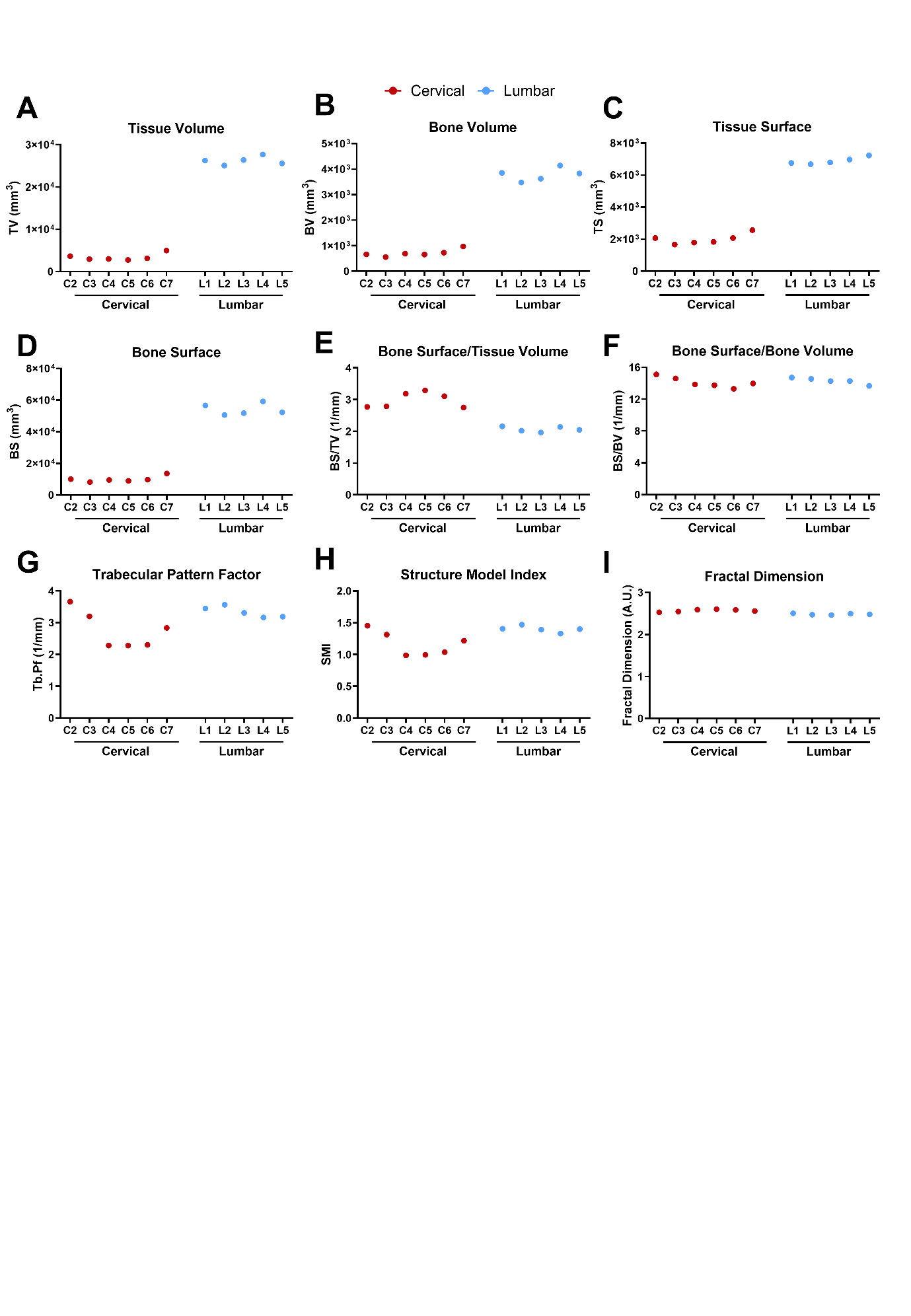

**Figure S2:** Variability in trabecular morphometric parameters across the cervical and lumbar spine. Quantification of three-dimensional vertebral trabecular tissue volume (A), bone volume (B), tissue surface (C), bone surface (D), bone surface/tissue volume (E), bone surface/bone volume (F), trabecular pattern factor (G), structure model index (H) and fractal dimension (I) from overview HiP-CT scans of the intact human spine. Tissue and bone volume along with surface measures were greater in the lumbar than cervical spine of this specimen, whereas bone surface/tissue volume was greater in the cervical spine. Trabecular pattern factor and structure model index showed modest level-specific variation, while fractal dimension remained relatively consistent across cervical and lumbar spine levels.

**References**

1. Walsh CL, Tafforeau P, Wagner WL, Jafree DJ, Bellier A, Werlein C, Kühnel MP, Boller E, Walker-Samuel S, Robertus JL, Long DA, Jacob J, Marussi S, Brown E, Holroyd N, Jonigk DD, Ackermann M, Lee PD. Imaging intact human organs with local resolution of cellular structures using hierarchical phase-contrast tomography. Nat Methods 2021;18(12):1532-1541. doi: 10.1038/s41592-021-01317-x

2. Brunet J, Walsh CL, Wagner WL, Bellier A, Werlein C, Marussi S, Jonigk DD, Verleden SE, Ackermann M, Lee PD, Tafforeau P. Preparation of large biological samples for high-resolution, hierarchical, synchrotron phase-contrast tomography with multimodal imaging compatibility. Nat Protoc 2023;18(5):1441-1461. doi: 10.1038/s41596-023-00804-z

3. Kyrieleis A, Ibison M, Titarenko V, Withers PJ. Image stitching strategies for tomographic imaging of large objects at high resolution at synchrotron sources. Nuclear Instruments and Methods in Physics Research Section A: Accelerators, Spectrometers, Detectors and Associated Equipment 2009;607(3):677-684. doi: 10.1016/j.nima.2009.06.030
